# Individualized Spatial Network Predictions Using Siamese Convolutional Neural Networks: A Resting-State fMRI Study of over 11,000 Unaffected Individuals

**DOI:** 10.1101/2021.03.22.436403

**Authors:** Reihaneh Hassanzadeh, Rogers F. Silva, Anees Abrol, Mustafa Salman, Anna Bonkhoff, Yuhui Du, Zening Fu, Thomas DeRamus, Eswar Damaraju, Bradley Baker, Vince D. Calhoun

## Abstract

Individuals can be characterized in a population according to their brain measurements and activity, given the inter-subject variability in brain anatomy, structure-function relationships, or life experience. Many neuroimaging studies have demonstrated the potential of functional network connectivity patterns estimated from resting functional magnetic resonance imaging (fMRI) to discriminate groups and predict information about individual subjects. However, the predictive signal present in the spatial heterogeneity of brain connectivity networks is yet to be extensively studied. In this study, we investigate, for the first time, the use of pairwise-relationships between resting-state independent *spatial maps* to characterize individuals. To do this, we develop a deep Siamese framework comprising three-dimensional convolution neural networks for contrastive learning based on individual-level spatial maps estimated via a fully automated fMRI independent component analysis approach. The proposed framework evaluates whether pairs of spatial networks (e.g., visual network and auditory network) are capable of subject identification and assesses the spatial variability in different network pairs’ predictive power in an extensive whole-brain analysis. Our analysis on nearly 12,000 unaffected individuals from the UK Biobank study demonstrates that the proposed approach can discriminate subjects with an accuracy of up to 88% for a single network pair on the test set (best model, after several runs), and 82% average accuracy at the subcortical domain level, notably the highest average domain level accuracy attained. Further investigation of our network’s learned features revealed a higher spatial variability in predictive accuracy among younger brains and significantly higher discriminative power among males. In sum, the relationship among spatial networks appears to be both informative and discriminative of individuals and should be studied further as putative brain-based biomarkers.

## 1. Introduction

Studies have investigated variability in human brain structure via standard assessment measures such as cortical thickness, sulcal depth, and cortical folding across individuals [1,2,3,4]. Later studies discovered the existence of relatively unique patterns in the human brain’s functional organization. In particular, significant variability in functional connectivity patterns (i.e., temporal dependence among brain network timecourses) between groups of subjects (e.g., controls and patients) has been reported in the past decade [5,6,7]. Other studies have shown that functional connectivity patterns can be used to predict individual traits [8]. Despite these findings, it is still unclear whether spatial relationships among functional networks in the brain can be linked to individuals uniquely. Motivated by that, we investigate the spatial maps of functional networks in this study and evaluate if the relationship between any two network spatial maps can predict whether or not the maps come from the same subject.

Brain neural activity can be indirectly recorded by functional magnetic resonance imaging (fMRI) based on the ensuing fluctuations in blood oxygenation level dependent (BOLD) signals [9,10]. Whole-brain resting-state fMRI (rs-fMRI) measures BOLD fluctuations while an individual is at rest, i.e., subjects are not performing an explicit task [11]. There has been significant interest in resting-state fMRI due to its lower design complexity and ease of acquisition (relative to task fMRI, where an individual needs to be able to perform a certain task) [12]. This advantage of rs-fMRI imaging renders it a widely used technology leading to many studies, including those of patients with particular conditions (e.g., comatose individuals or Alzheimer’s patients) [13]. Multiple studies have also shown rs-fMRI data can be used to estimate subject-level differences [14]. Various model- and data-driven methods (such as seed-based correlation analysis [15], independent component analysis (ICA) [16,17], graph methods [18], and clustering algorithms [19]) analyze whole-brain rs-fMRI scans to identify spatially distinct but functionally correlated regions, also called resting-state networks (RSNs). Compared to other methods, ICA can identify maximally statistically independent RSNs with less prior information. Also, it is able to capture artifacts and noise while separating these from the RSNs [9]. These advantages have led to the widespread use of ICA to analyze rs-fMRI data.

Studies that utilize ICA for processing rs-fMRI data often analyze temporal patterns of brain activity, such as the pairwise correlation between RSN-specific timecourses [20,21,22,7]. Despite the predominance of studies on temporal characteristics of rs-fMRI data, the spatial characteristics of functional networks also carry remarkable information and possess distinctive patterns for characterizing subjects [23,24,25]. Recent studies have reported that ICA patterns can be used to classify group membership (e.g., patients versus controls) [26]. In this article, we shed more light on a newer and yet more challenging problem, namely, the discrimination of subjects solely from their underlying spatial map networks by learning inter-network relationships in a verifiable way. To the best of our knowledge, this problem has not been studied before, yet it has important implications for future studies of brain-based biomarkers and distributed networks.

The majority of studies that analyze resting-state connectivity rely on comparing functional patterns between groups of people in some way. Some of these studies have tried to distinguish unaffected control subjects from symptomatic ones, such as those with Alzheimer’s disease [7,27], schizophrenia [8,28,29], language-impairment [5], and autism [30,31]. In comparison, others have provided models to classify subjects according to their sex [32,33] or age [34] using functional connectivity. Altogether, although these studies attain individual-level predictions, they are ultimately focused on the use of group-level (dis)similarity. However, as mentioned above, there has been much less focus on individual-level *spatial* heterogeneity in brain functional interactions.

A few studies have shifted the focus to link such patterns to individuals [6,14,35,8]. More specifically, Finn and colleagues [6] identify subjects based on their whole-brain *and* network-based functional connectivity. They scanned the brain of 126 subjects over six sessions when performing working memory, motor, language, and emotion tasks as well as at resting state. Then, for each subject, the functional connectivity derived from each session was compared (using Pearson correlation) to the set of functional connectivity of all subjects from other sessions. Another study [14] computed the variability in the intrinsic functional connectivity of a small rs-fMRI dataset of 23 healthy subjects collected over six months with five scan sessions per individual. The individual variability in some brain regions was evaluated, including frontal, temporal, and parietal lobes, and shown to be higher than in other regions. In a quite different approach, a linear model was proposed in [35] for the prediction of a region of interest (ROI) time series from another ROI time series. A model was fit to assess if the prediction is unique for each subject, i.e., the prediction pattern was distinctive between 27 individuals. Such models suffer from a few fundamental shortcomings. First, they usually require multiple scans from each subject for proper training, which is costly and time-consuming; for instance, it took over six months for the authors in [14] to complete the data acquisition phase. Second, due to the lack of high-sample-sized datasets for such longitudinal models, it is hard, if not impossible, to train the high-capacity models that are typically better at capturing complex features and, thus, such models are not otherwise generalizable due to overfitting [36]. Furthermore, all these studies disregard the spatial maps of functional brain networks in their analysis, while several studies have shown such information can be used as biomarkers to characterize individuals [37,38,39].

Here we use a large resting-state fMRI dataset to identify discriminative features of brain activity between unaffected individuals to address the shortcomings above. Our goal is to characterize individuals based on high-level features learned from their functional brain network spatial patterns and from pair-wise comparisons between subjects given a pre-selected network pair. We are also interested in observing which specific networks are more informative for such task. Accordingly, we develop a deep neural network-based framework that can detect subjects based on high-level difference features in the spatial patterns of their functional networks, when those networks are different (e.g., auditory and visual networks) for each subject. Our assumption is that there exists some high-level spatial pattern that underlies all such functional networks but is unique to each person. Thus, we hypothesize that subjects can be differentiated from each other by contrasting such a pattern if it can be extracted and represented in a latent feature space where the distances make biological sense. In other words, our aim is to learn a functional brain pattern that 1) is unique to the individual brain and 2) can be mapped to a unified feature space where differences in subject labels translate, by design, to L1 distances. In that respect, it does not matter which network of the brain we are selecting from an individual as long as we can infer the latent feature(s) from it, which can then be used to capture differences. Given the possibly nuanced complexity and unpredictable nature of such spatial (dis)similarity features, it is fair to conjecture they may be successfully learned via deep neural network model representations.

Our approach of characterizing brain samples using a pair of functional brain networks is different from the few previous studies in several ways. First, we develop a framework to capture complex functional patterns in the brain and do so in a region agnostic manner (i.e., regardless of the selected brain network pair). Second, we transform the problem of subject identification into subject comparison by taking advantage of the Siamese architecture [40,41], thus reducing the original multiclass problem into a binary classification problem (class 1 (or positive) if two input spatial maps are from the same subject, class 2 (or negative) otherwise), which is moderately easier to train. Third, we leverage the nature of this task to obtain a relatively large augmented dataset of *pairs of subjects*, effectively boosting the predictive power of our end-to-end trained models. Furthermore, our models do not require multiple scans per subject, which is a limitation in some previous studies. Finally, as will be discussed, each trained model works on a preselected pair of functional networks and, thus, can capture relationships between brain regions, contrary to previous work that is limited in this regard.

This paper is organized as follows. In section 2, we describe the data and the procedure underlying data collection and preprocessing and subsequently introduce our model in more detail. In section 3, we evaluate our model on a held-out test set using the Monte Carlo cross-validation approach and shed more light on age and sex differences in its performance. Section 4 reflects on our observations and the model’s performance. Finally, section 5 concludes our paper and suggests future directions to continue this line of research.

## 2. Materials & Methods

### 2.1. Participants

We retrieved the resting-state fMRI dataset from the UK Biobank [42]. At the time of retrieval, this dataset included 19831 subjects, out of which 13668 were self-reported as healthy (unaffected) adult participants. The subject fMRI scans underwent quality control, and subjects were excluded if the scans met the following criteria: marked as unusable by UK Biobank, visual inspection of mean maps for gross anomalies, absolute framewise displacement (FD) higher than 0.3mm, Matthews correlation coefficient (MCC) between the binarized study-specific mask and the subject mask lower than 0.8, and failure to complete ICA estimation. 11754 subjects were finally retained for the analysis after quality control, with the included participants’ ages ranging from 45 to 80 (62.56 ± 7.38) years. The dataset was well balanced in terms of the participants’ sex (c.f. Table 1).

**Table 1.** Subjects’ demographics.

|  | **Population Number** | **Age (years)** |  |  |  |  |  |  |
| --- | --- | --- | --- | --- | --- | --- | --- | --- |
|  |  | *Mean* | *SD* | *Min.* | *25%* | *50%* | *75%* | *Max.* |
| **All** | 11754 (100%) | 62.56 | 7.38 | 45 | 57 | 63 | 68 | 80 |
| **Male** | 5772 (49%) | 63.08 | 7.54 | 45 | 57 | 64 | 69 | 80 |
| **Female** | 5982 (51%) | 62.07 | 7.19 | 46 | 56 | 62 | 68 | 80 |

### 2.2. Data Acquisition & Preprocessing

All participants were scanned once by a 3-Tesla (3T) Siemens Skyra scanner with a 32-channel receive head coil, all acquired in one site. A gradient-echo echo planar imaging (GE-EPI) paradigm was used to obtain resting-state fMRI scans. The EPI-based acquisition parameters include multiband acceleration factor of 8 (i.e., eight slices were acquired simultaneously), no iPAT, fat saturation, flip angle (FA) = 52°, spatial resolution = 2.4 × 2.4 × 2.4mm, field - of - view (FOV) = (88 × 88 × 64 matrix), repeat time (TR) = 0.735s, echo time (TE) = 39ms, and 490 volumes. Subjects were instructed to stare at a crosshair passively and remain relaxed, not thinking about anything, during the six-minute and ten-second resting-state scanning period.

The preprocessing steps performed by UK Biobank are as follows. An intra-modal motion correction tool, MCFLIRT [43], was applied to minimize the distortions due to head motion. Grand-mean intensity normalization was used to scale the entire 4D dataset by a single multiplicative factor to compare brain scans between subjects. The data were filtered by a high-pass temporal filter (Gaussian-weighted least-squares straight-line fitting, with σ = 50.0 s) to remove residual temporal drifts. Geometric distortions of EPI scans were corrected by using the FSL’s Topup tool [44]. EPI unwarping is followed by a gradient distortion correction (GDC) unwarping phase. Finally, structured artefacts are removed by ICA+FIX processing (Independent Component Analysis followed by FMRIB’s ICA-based X-noiseifier [45,46,47], with no lowpass temporal or spatial filtering up to this point. More details on the UK Biobank imaging protocol and preprocessing steps can be found in [42]. In addition, the data were then normalized to an MNI EPI template using FLIRT followed by SPM12, old normalization module. Finally, the data were smoothed using a Gaussian kernel with FWHM = 6mm.

### 2.3. Group Independent Component Analysis

We applied fully automated spatially constrained ICA using the NeuroMark approach [48] on the 4D preprocessed UK Biobank rs-fMRI data from Section 2.2. In the Neuromark approach, a template of replicable independent components (ICs) was constructed after spatially matching correlated group-level ICs between two healthy control fMRI datasets - genomics superstruct project (GSP) and human connectome project (HCP). The estimated network template was then used as a prior for a spatially constrained ICA algorithm applied to each UK Biobank subject individually. This identified 53 functionally relevant resting-state networks (RSNs) for each individual that are maximally spatially independent (see Fig. S1 and Table S1). Each subject-specific RSN is represented by a spatial map of size 53×63×52 voxels and its associated time course of 490-time points. RSNs are grouped into seven domains, namely subcortical (SC), auditory (AU), sensory-motor (SM), visual (VI), cognitive control (CC), default mode (DM), and cerebellar (CB), by functional similarity (see Fig. 1). This work used the subject-specific spatial maps (henceforth referred to as networks) as input to our model and a basis for all subsequent analyses.

**Fig. 1.**
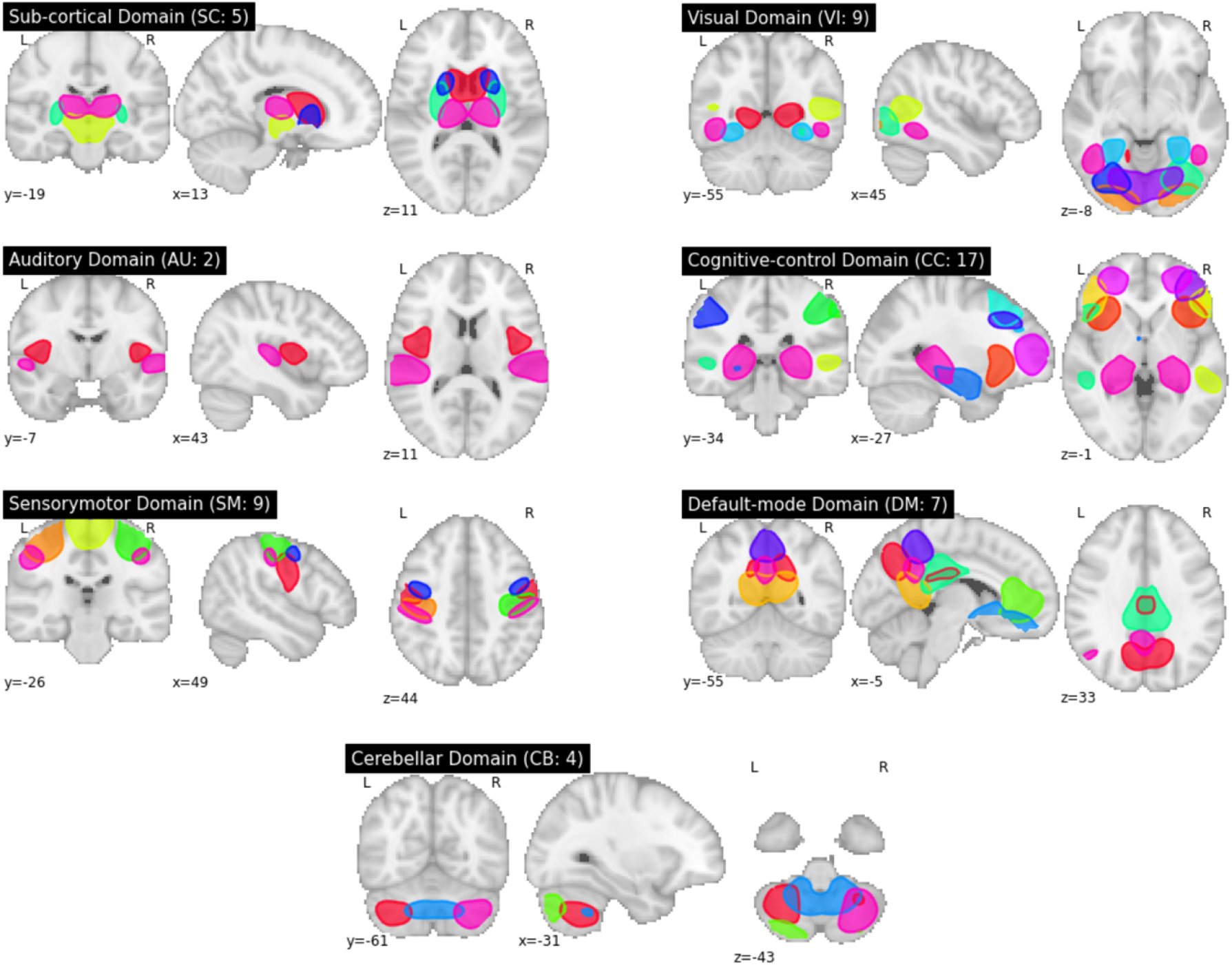
Group ICA-derived spatial maps. Spatial maps are grouped into seven domains, subcortical (SC), auditory (AU), sensory-motor (SM), visual (VI), cognitive control (CC), default mode (DM), and cerebellar (CB), each of which contains 5, 2, 9, 9, 17, 7, and 4 networks, respectively.

### 2.4. Generating Input Pairs

We split our preprocessed data *by subjects* into training, validation, and test sets with a proportion of 60%, 20%, and 20%, respectively (see Table S2-4 for the statistics of subjects in each set). The number of *same*-subject pairs created for each set was equal to the total number of subjects in each set, and all these pairs were labeled ‘class 1.’ Likewise, an equal number of *different-subjects* pairs (labeled ‘class 2’) were created for each set by randomly selecting subjects from the corresponding set (for a total of *N* pairs out of 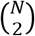 possible pairs, where *N* is the total number of subjects in each set). As discussed before, input pairs are comprised of two networks instead of one. This is important as, otherwise, instead of learning patterns that govern the relationships within networks, the model gradually (i.e., within the course of training) tends to learn to make an elementwise comparison of the input voxels. We trained one model for each pair of the 53 functional networks using the TReNDS high-performance computing GPU cluster. In total, 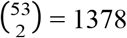 models were trained.

### 2.5. Model Architecture

We provide an end-to-end trainable deep learning model for learning a non-linear dissimilarity metric based on patterns of spatial relationships in functional networks that characterize each subject. We use balanced (i.e., same-subject vs. different-subject) preprocessed input-pairs for training, and the objective is to adjust network weights such that input pairs that belong to the same subject produce a low dis-similarity score whereas those pertaining to different subjects generate a large score. Our proposed framework comprises two major building blocks, one based on the convolutional neural networks (CNNs), which are superior in capturing high-level features from imaging data modalities [49], and the other based on a Siamese network that can efficiently generate features for differentiating inputs [40,41]. In the following sub-sections, we shed more light on the architecture of each module.

### 2.6. Convolutional Neural Networks (CNNs)

2D CNNs and, more recently, 3D CNNs have gained attention in various domains that involve image processing. CNNs can pick up high-level and yet subtle, task-specific features if trained on task-appropriate loss functions. For our analysis, the desired set of features stem from patterns of functional brain relationships that can help characterize subjects, yet in a relatively network-agnostic way. In other words, these features should be comparable between different subjects, even if they are derived from different spatial maps. This is indeed a crucial component of our design since using spatial maps from the same networks drives the training course towards a point in the parameter space that corresponds to a voxel-to-voxel verbatim comparison. In other words, instead of learning a pattern that characterizes brain activities, the model will tend to see if the two images are identical voxel-wise. We therefore prevent this by utilizing network spatial maps as input and two CNN-based child networks that can generate informative features linked to each other via a Siamese architecture, as will be discussed shortly.

Fig. 2 depicts the architecture of the employed 3D ConvNet that extracts features from a supplied spatial map with a resolution of 53×63×52 voxels. We used three convolutional layers with 16, 32, and 64 filters, respectively, and used kernels of 3×3×3 for each of the layers. We used ReLU activations for both convolutional and later fully connected layers. Furthermore, we augmented the convolutional layers with max-pooling layers (with a kernel size of 2×2×2 and stride of 2) to reduce the feature maps dimensionalities. The features in the last feature map are flattened and then fed into a single fully-connected layer with a ReLU of size 128 neurons. We used 3D and 1D batch-normalization as a regularization for all layers. This is especially important due to the large dimensionality and complexity of the model. The DL *encodings* (i.e., final feature values) serve as a summary of the input 3D spatial map, capturing the non-linear pattern of functional relationships underlying the input brain network pair when trained based on a loss that quantitatively factors in our goal, as discussed in the next section.

**Fig. 2.**
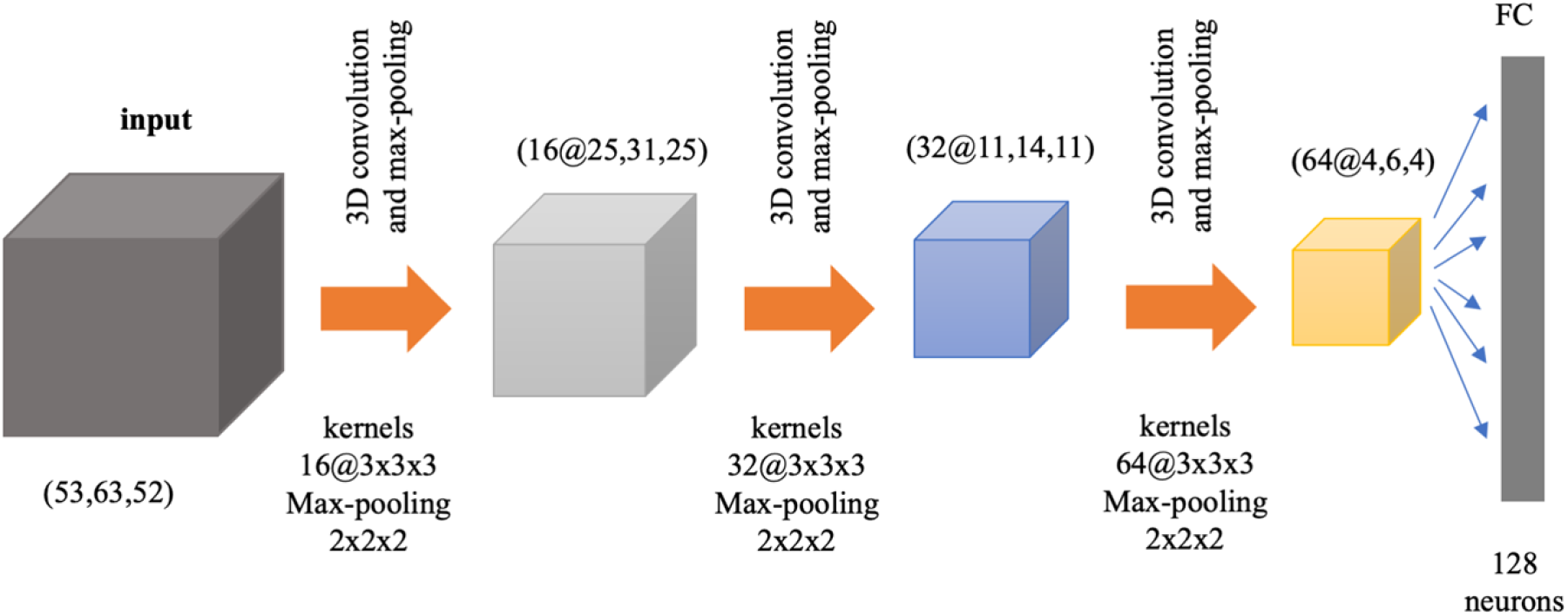
CNNs architecture. The CNNs block consists of three 3D convolutional layers, with kernels of sizes 3×3×3, each of which is followed by a max-pooling layer with a kernel of size 2×2×2.

### 2.7. Siamese Architecture

For our brain-network classification task, we use a modified Siamese-based CNN model, seeking to map input image pairs to a shared target space in which both spatial map images are comparable. In other words, we learn separate mappings that transform the (spatial) input spaces of two *different* functional brain network maps into a shared high-level space. In that space, the distance between subjects (irrespective of the brain network region) represents how closely related their corresponding functional networks are. By training specifically on *different* brain network pairs, we ensure that closely related network representations learned by the model are not merely driven by spatial similarity but, rather, by having their origin on the same subject. Fig. 3 shows the block diagram of our proposed model. It is comprised of two CNN-based *child* networks with identical architectures (referred to as sub-networks hereon), but not sharing weights. Each sub-network takes a different functional brain map network image, namely *X*_1_ and *X*_2_ (e.g., visual network and auditory network), to generate consistent and comparable representations *R*_1_(*X*_2_) and *R*_2_(*X*_2_), respectively. Then, optimizing for a Siamese-based architecture using the binary cross-entropy loss drives parameters of the CNN sub-networks towards a point in the parameter search space where projections lie in distant locations when subjects are different. Our experimentation suggests that this is more easily attained with independent, rather than tied, weights for each child network. This choice is also driven by the problem at hand. Using shared weights is a common approach if the objective is to learn a generic representation for both input data modes (here, brain network spatial maps), whereas using independent weights is more suitable if learning mode-specific representations instead. Thus, the use of separate weights is more appropriate for our objective of training models that learn if two modes (i.e., brain network spatial maps) correspond to the same subject or not. Additionally, we assume that a softmax-normalized L1 distance, trained through the binary cross-entropy loss, will induce both sub-networks to converge into a shared feature space.

**Fig. 3.**
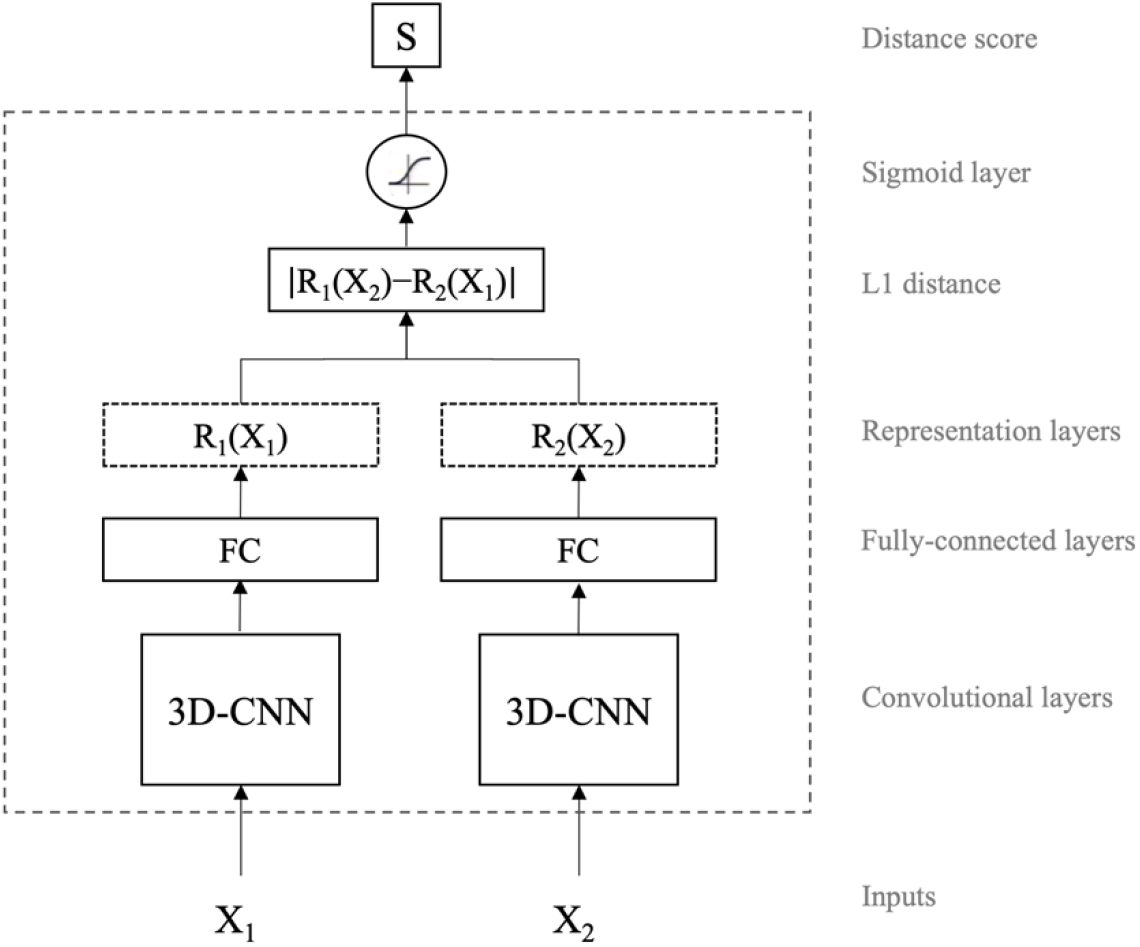
High-level architecture of the proposed model. Siamese networks take two images, X_1_ and X_2_, and learn a distance function between the two vector representations of the input images. The model outputs distance score close to one if the two inputs are from two different objects (in this case, different subjects) and close to zero otherwise. The weights of the CNN child networks are updated independently and are not tied/shared.

We train this network for 200 epochs using stochastic gradient descent with a binary cross-entropy loss and a learning rate of 0.001. The model is given by *loss* = −*t*log(*S*) — (1 — *t*)log(1 — *S*) where *t* is the true label, *S* = *sigmoid*(∑ *W_i_d_i_*), *d_i_* = |*R*_1_(*X*_2*i*_) — −*R*_2_(*X*_1*i*_)|, and *i* is the dimensionality index of the feature space. The training is stopped using the early-stopping strategy of no improvement in validation accuracy after 14 consecutive epochs. Furthermore, we repeat the training course three times and select the best validation model (i.e., the one with the highest validation accuracy) for testing on the held-out test data.

## 3. Experiments

In this section, we evaluate the performance of the proposed model when trained on different pairs of functional brain networks, on unseen (held-out) test datasets. In section 3.1, we compare and contrast the accuracy achieved when we select networks from different domains. In addition, we discuss and analyze the prediction performances across three held-out test set sample cohorts according to the class labels we attempt to predict: network pairs from the same subjects, network pairs from different subjects, and the entire (aggregate) set of all network pairs (same and different subject pair samples together). Using these cohorts enables us to evaluate our model architecture’s performance under different scenarios and provides insight into brain function. In sections 3.2 and 3.3, we analyze the relationship between cognitive features, age and sex, and model performance.

### 3.1. Analysis of Performance in Different Functional Network Domains

To investigate potential variation in model performance by network pairs, we evaluated the proposed model on the entire held-out test set (i.e., the cohort containing all subject pair samples), which is comprised of an equal number of same-subject and different-subject network-pair samples. Fig. 4A shows a heatmap of the mean model accuracies between domains (see the fine-grained brain network connection-level accuracies in Fig. S2). According to the heatmap, there exist spatial network relationship patterns with strong prediction capability stemming from SC-SC, SC-CC, SC-DM, SC-VI, VI-VI, and DM-DM domain-pairs. On the other hand, network pairs including an auditory network appear to contain less discriminative (yet still significant) features. Overall, it turns out that the amount of discriminative information that can be captured from a given network pair is at least partly domain-dependent (see Table S5 for a two-sided two-sample *t*-test of the mean prediction accuracy between domains and the corresponding *p*-values). Furthermore, to assess the reproducibility and generalizability of the results, we repeat our training-testing approach using the Monte Carlo cross-validation (MCCV) approach with five repeats (the held-out test set changes for each repeat). Fig 4B shows a boxplot of the corresponding accuracies for each pair of networks that fall under SC-SC and AU-AU domain pairs (i.e., the highest and the lowest performant domain pairs according to Fig. 4A, respectively). From the figure, it is evident that our results are reproducible up to a negligible variation in model performance.

**Fig. 4.**
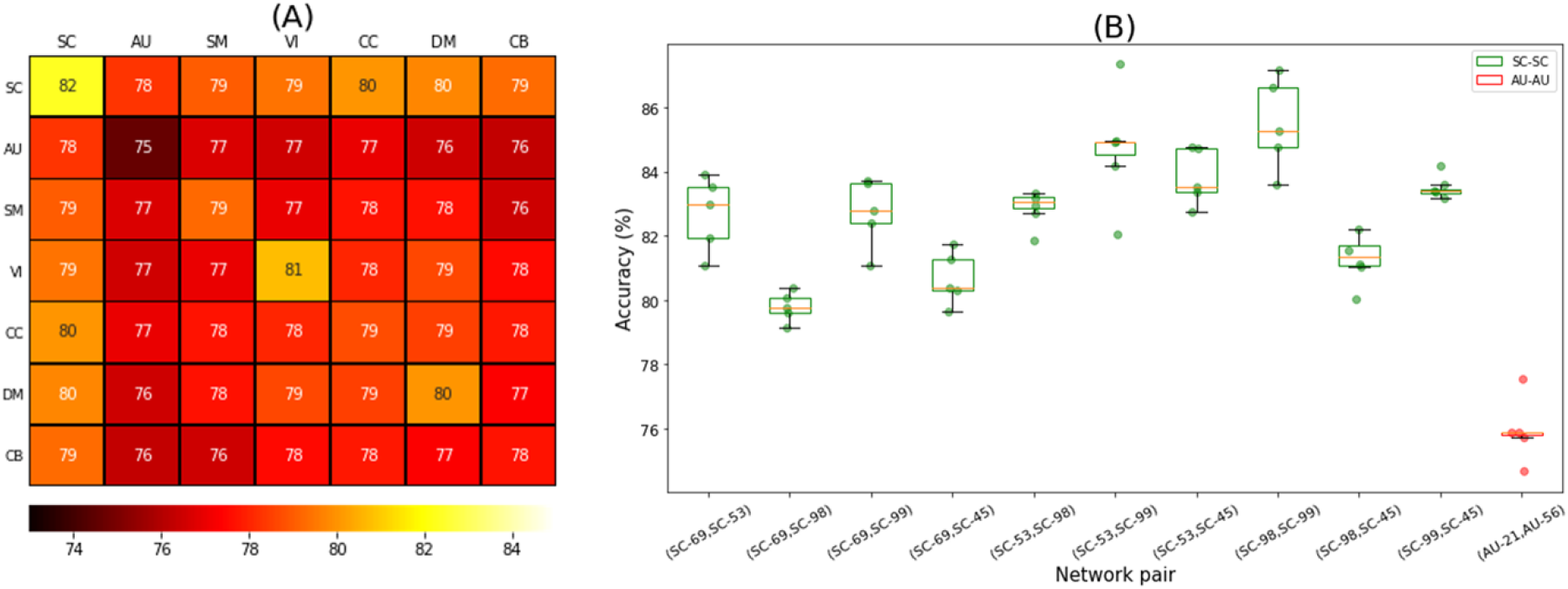
(A) Mean domain accuracy heatmap. Each cell shows the mean accuracies of all models trained on spatial networks belonging to the indicated domain-domain pair. The resulting mean accuracies show that domains have different predictive power for characterization of subjects based on brain activity. A comparison of SC to the other domains reveals that its underlying networks yield the most discriminative features. On the other hand, the AU domain appears to produce the least discriminative features relative to other regions in the brain. **(B) Boxplot for the accuracies derived from the MCCV evaluation.** The results show low variation across the five repeats for pairs of networks that fall under SC-SC and AU-AU domain pairs, which provides supporting evidence for the generalizability of the proposed model. Green and red boxes represent SC and AU network pairs, respectively. The dots are the results of MCCV for each network pair. The X-axis indicates the Neuromark network ids.

We were also interested in investigating whether any network pairs were more feature-rich for either of the same-vs. different-subject cohorts in the classification task. This is an important question as significant differences can serve to support recommendation guidelines for optimally choosing certain network pairs for different goals. In light of that, we experimented by partitioning the test samples into a cohort of network pairs from the same subjects and a cohort of pairs with different subjects. Figures 5A and 5B depict the spatial connectograms of the top 2% network pairs with the highest sensitivity (we treat same-subject samples as the positive class) and specificity, respectively (see Fig. S3A and S3B for the full results). We grouped connectograms based on their functional domains, i.e., SC, AU, SM, VI, CC, DM, and CB, which contain *n* = 5, 2, 9, 9, 17, 7, and 4 networks, respectively. Specifically, Fig. 5A shows that the highest sensitivities are mostly from brain regions in the subcortical and cognitive control domains. While according to Fig. 5B, brain regions in the visual domain exhibit the highest specificities. Thus, comparing the two connectograms suggests that different domains offer distinct patterns of contribution towards identifying same-vs. different-subject pairs.

**Fig. 5.**
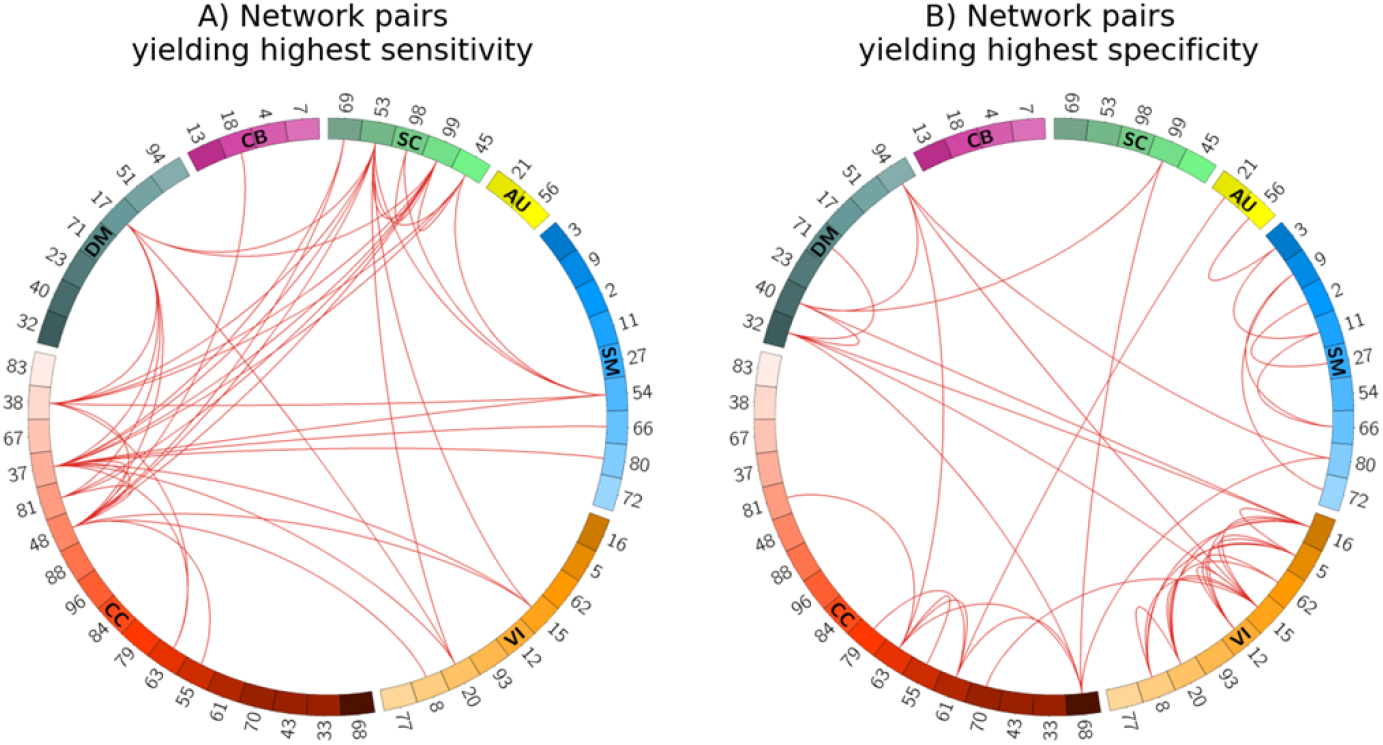
Spatial connectogram of sensitivity (A) and specificity (B) highlighting the top 2% network pairs. Sensitivity and specificity show the percentage of network pair samples correctly identified as the same- or different-subject cohorts, respectively. Fig. 5A suggests that networks from the subcortical (green) and cognitive control (red) domains are among the most useful for characterizing same-subject samples, both intra- and interdomain. On the other hand, the high specificity of VI-VI network pairs (orange) in Fig. 5B indicates better characterization of different-subject samples. The numbers shown on the connectograms represent the specific Neuromark RSN network indices (See Table S1 for a list of the network names). The connectograms were generated using the Circos tool [50].

### 3.2. Impact of Sex on Performance

We also evaluated the role of sex on the prediction performance separately for the same- and different-subject cohorts. First, we considered the same-subject cohort and compared male vs. female subcohorts within it. We observed that the model’s sensitivity when two networks are selected from male subjects is larger than when they are selected from female subjects, which was the case for 71% of network pairs (See Fig. S4). This suggests that male functional networks have more uniquely identifying individual patterns, especially in the cerebellar domain, compared to females. This finding is consistent with studies of brain structure that maintain males’ brains have more variability than females [51,52]. Indeed, higher variability in brain structure is conducive to the existence of more distinct patterns in males that help characterize male subjects’ identities more accurately.

To better visualize the difference in performance, we subtracted the female cohort’s sensitivity scores from that of males for each network pair and selected the network pairs corresponding to the highest (positive) and lowest (negative) 1% score difference, as illustrated in Fig. 6A. According to the figure, from the pairs in which males have higher sensitivity (i.e., the red arcs), the superior parietal lobule (SPL) network (indicated as SM27) shows more outgoing arcs in the connectogram. This may suggest that the SPL network activity varies more (or, is more uniquely identified) in males than females. To the best of our knowledge, this is the first time such a pattern has been identified. Studies have shown that the SPL network is linked to spatial processing tasks, especially in mental rotation [53,54]. Another interesting research has reported stronger activity of these networks among males [55]. Altogether, these observations seem to generally indicate that while men might do better at spatial orientation tasks than women, there is more variation to their brain functional engagement in this task than women. Hence, the significance of SM27 as a functional brain activity biomarker for classification of males from females should be considered. Likewise, among networks with the lowest (most negative) sensitivity difference between males and females (i.e., the blue arcs in Fig. 6A), CC70 presents the most outgoing arcs, suggesting this network is more discriminative in females.

**Fig. 6.**
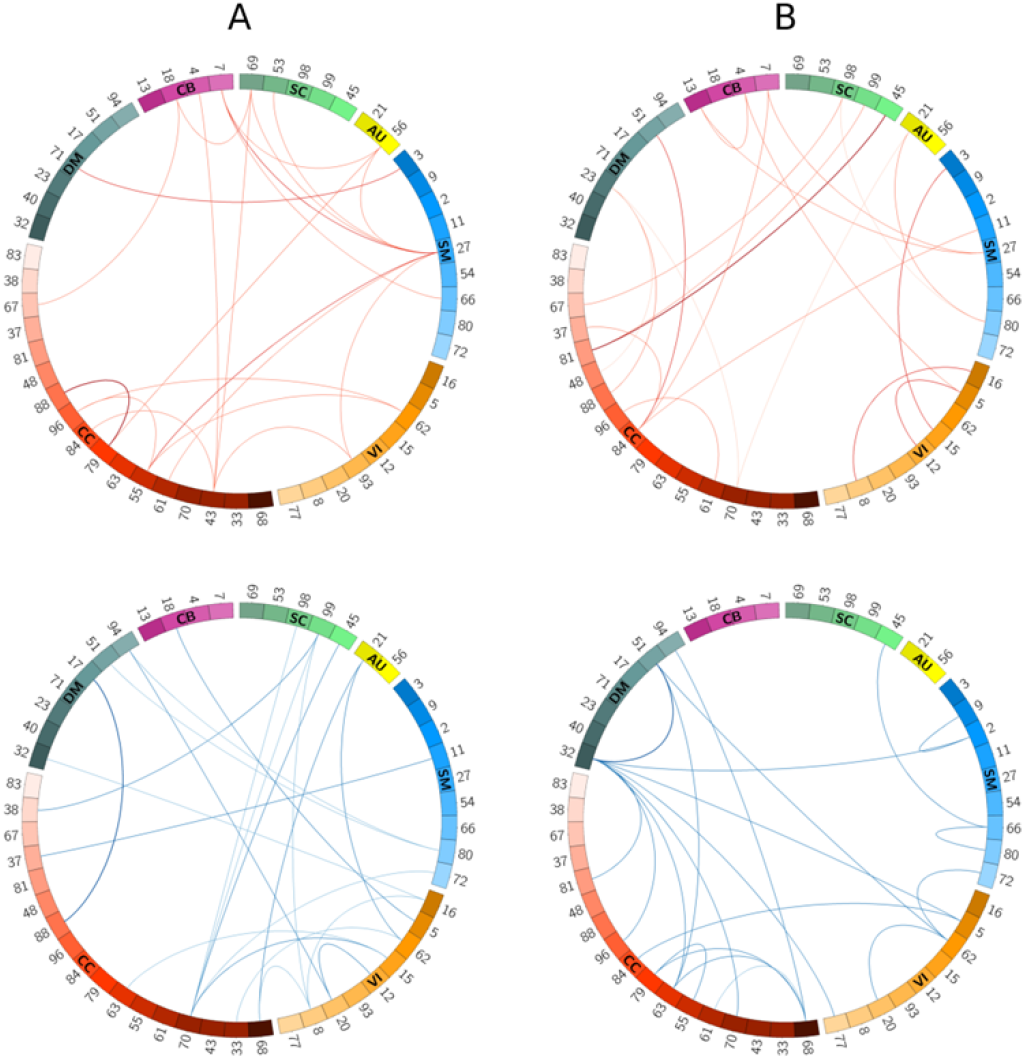
Spatial connectogram of difference in sensitivity (A) as well as the difference in specificity (B) between sex-based groups highlighting the top 1% positive (top row) and negative (bottom row) differences. Fig. 6A: scores are calculated by subtracting sensitivities of the female subcohort from the male subcohort. SM27, when paired with other domains, produces higher specificity in males, while CC70 yields higher specificity in females. Fig. 6B: scores are calculated by subtracting specificities of different-sex from same-sex subcohorts. The SM and CB networks produce higher sensitivities for different-sex samples, whereas the CC networks predict better in the same-sex subcohort. Blue and red color lines represent the two ends of the difference spectrum. Numbers represent indices for the ICA-derived Neuromark RSNs. The connectograms were generated using the Circos tool [50].

Finally, we considered the set of different-subject samples. This set naturally splits into same-sex (male-male or female-female pairings) and different-sex (male-female pairings) subcohorts, for which we conducted assessments analog to those shown in the male vs. female discussion above. Interestingly, we observed higher specificity in the different-sex cohort for 86% of network pairs. According to the results of this assessment, the model performs better (higher specificity) when samples belong to different sexes, for most network pairs. This is likely because our model is capable of learning sex-rich features from the network spatial map patterns, even though sex has not been used as an input to our model. The corresponding highest and lowest 1% specificity differences are visualized in Fig. 6B. For the most negative differences (same-sex specificity lower than for different-sex), we observed that network pairs including either the default mode (especially DM32 and DM51) or the cognitive control networks had the most outgoing links, suggesting more unique variations in the different-sex cohort. For those network pairs with positive specificity differences, on the other hand, the CC networks, especially CC84, appear more frequently in the connectogram, suggesting that more unique patterns occur in the same-sex cohort when network pairs include a CC network.

Furthermore, we assessed the previous experiments’ results from a statistical perspective (see Table S6). We performed a two-sided two-sample *t*-test which shows the significance of the mean sensitivity or specificity difference between the two sub-cohorts of each sex-based assessment. Accordingly, we observed that the mean sensitivity in the male sub-cohort is significantly different in the female sub-cohort, and the mean specificity in the different-sex sub-cohort is significantly different from the same-sex sub-cohort. These seem especially true for the cognitive control domain.

### 3.3. Impact of Age on Performance

Next, we analyzed the relationship between age and model performance. Similar to experiments in the previous section, we divided the test set into same- and different-subject cohorts. We computed the sensitivity of subjects of ages below 52 and above 72, which make cohorts of size 260 and 268 subjects, respectively. Our results revealed that the younger cohort’s prediction performance is significantly higher than that of the older cohort for 66% of network pairs (see Fig. S5A). This is especially the case for the cognitive control (*p* < 2*e* — 22, see Table S7), sensory-motor (*p* < 6*e* — 18), and default mode (*p* < 1*e* — 11). In a similar experiment, we computed the specificity for all pairs of networks within the different-subject cohort for 139 and 158 subjects with age below 57 and above 69 years old, respectively (see Fig. S5B). For most domains, the model performance for younger brains appear to be stronger than older ones, especially for the cognitive control (*p* < 4*e* — 8) and cerebellar (*p* < 2*e* — 8) domains.

To shed more light on the role of age on our model performance, we computed pairwise sensitivity and specificity differences between the young-old cohorts and then picked the highest and lowest 1% of the resulting scores. Fig. 7A and 7B show these differences for the sensitivity and specificity metrics, respectively. Based on these figures, from the pairs with the largest sensitivities (the arcs colored in red) networks CC61, CC79, and SC69 when paired with a number of other networks appear to be more discriminative. On the other hand, among the networks with the lowest sensitivity difference score (arcs colored in blue) networks CB18 and SC98 are linked with many outgoing arcs suggesting that these networks contain more unique patterns within the cohort of old subjects. Altogether, the aforementioned networks can be used in age-related tasks where we are interested in comparing intra-subject networks. Likewise, considering Fig. 6.B, networks from the CC and SM domains when linked with other networks (Fig 7B, bottom row) along with the (CC70, VI5) network pair (Fig 7B, top row) are well suited for age-related tasks using patterns of networks between subjects (e.g., classifying younger vs. older individuals).

**Fig. 7.**
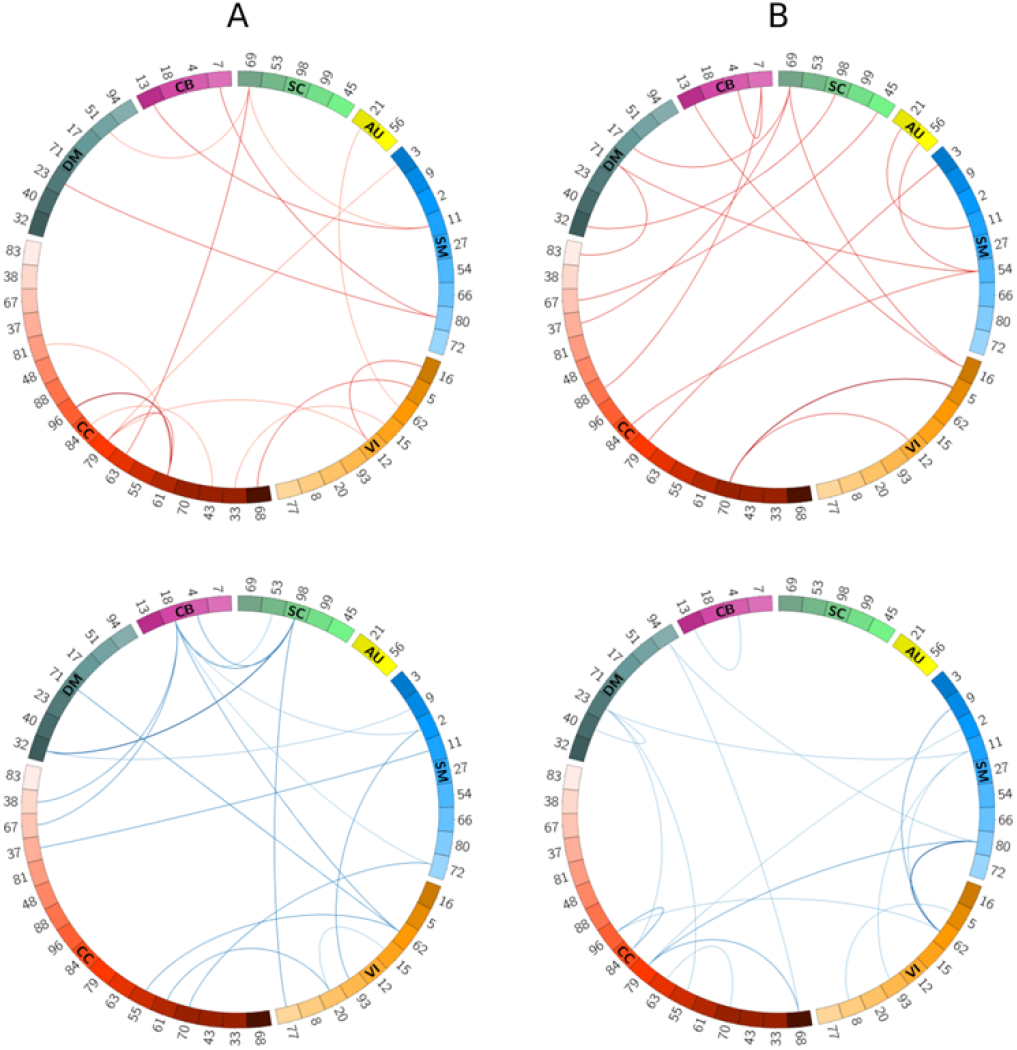
Spatial connectogram of difference in sensitivity (A) as well as the difference in specificity (B) between young and old groups with the highest (top row) and lowest (bottom row) 1% values. All the connectograms show the difference scores computed from deducting old subjects from young subjects. Fig. 7A suggests CC61 and CC79, and SC69 networks contain more discriminative features in young subjects. On the other hand, networks of CB18 and SC98 perform better when subjects are old. Fig. 7B shows the network pair CC70-VI5 has the largest specificity difference between young and old subjects. In contrast, the combination of the CC and SM networks can better predict the dis-similarity metric when subjects are older. The connectograms were generated using the Circos tool [50].

## 4. Conclusion

In this study, we showed for the first time that pairwise-relationships between ICA-based spatial maps can predict whether or not two networks belong to the same subject. As such, we proposed a Siamese-based model that extracts individualized patterns by pairwise comparison of spatial network maps (e.g., an auditory and a visual network). In our model, we combined a Siamese architecture with convolutional neural networks to learn high-level features describing 3D functional brain network maps suitable for the downstream comparison of subjects in our prediction task. Using the features extracted from the CNN networks, our model generated a non-linear distance metric representing the chance the two networks are from same or different subjects. Our results of training the proposed model for all possible network pairs showed that different pairs of functional networks contributed differently to the network prediction task. This was especially true when networks were selected from the subcortical domain (with the highest accuracy) or the auditory domain (with the lowest accuracy). The model’s high performance in subcortical networks suggested the superiority of such networks being used for future end-point tasks, such as age classification, brain fingerprinting, etc., under the proposed framework.

We also provided guidelines for neuroimaging-based prediction tasks by investigating which network pairs were more feature-rich in each of the same-subject and different-subject cohorts. We observed that networks from the subcortical and cognitive control domains, and especially their combinations, demonstrate more variability in the cohort of same-subject samples. Therefore, we suggest such pairs of networks are well suited for analyzing spatial interactions, for example, for predicting diseases. On the other hand, in the different-subject cohort, network pairs from the visual domain resulted in more accurate models and can later be used for a task that scrutinizes networks between subjects, e.g., for identifying same-sex samples.

Further analysis of our results revealed that the performance of our model depends on subjects’ age and sex. From the sex-based assessments, we observed in most cases (71%) the model performed better in males corroborating the previous studies’ findings, which showed that males’ brain networks have larger variability (i.e., more distinct patterns) than females’ [51,52]. Especially for the first time, we showed functional activity in the SPL network varies more within males than females. Future assessment and study of potential sex bias in these findings is warranted, and approaches such as [56] could be easily adapted to our framework for such purposes. Moreover, for 86% of network pairs, we observed higher specificity among different-sex subjects than same-sex subjects. This suggests higher discriminative patterns between networks of different sex compared to networks of the same sex. When it comes to the impact of age, our results showed that the prediction performance among the younger cohort is significantly higher than that of the older cohort for most network pairs. Overall, our age and sex-related findings showed that our model could learn sex and age-rich features from the spatial maps, even though sex and age have not been explicitly used as an input to our model.

Overall, despite the widespread use of timecourse-derived information (i.e., functional connectivity) in prediction-focused neuroimaging studies, spatial maps are feature-rich data sources that can serve as surrogates to timecourse data or even be used as complementary source input. As future work, we encourage to utilize the same approach to investigate whether similar patterns exist among diseased and healthy cases. Moreover, we hypothesize that our model can serve as the first step towards brain fingerprinting through discovering unique functional patterns that can characterize people in a unique way.

## Author Contributions

Reihaneh Hassanzadeh proposed and implemented the main idea, trained models, analyzed the results, and drafted the paper. Rogers F. Silva, Yuhui Du, Mustafa Salman, Anna Bonkhoff, Zening Fu, Thomas DeRamus, Eswar Damaraju, Bradley Baker, and Anees Abrol preprocessed the data, performed quality checks, and extracted the ICA components. Vince D Calhoun served as the PI and our guide throughout the project. All authors provided input and edits to the manuscript.

